# Accurate allele frequencies from ultra-low coverage pool-seq samples in evolve-and-resequence experiments

**DOI:** 10.1101/244004

**Authors:** Susanne Tilk, Alan Bergland, Aaron Goodman, Paul Schmidt, Dmitri Petrov, Sharon Greenblum

**Affiliations:** Department of Biology, Stanford University, Stanford CA 94305; Department of Biology, University of Virginia, Charlottesville VA 22904; Department of Biology, University of Pennsylvania, Philadelphia PA 19104

## Abstract

Evolve-and-resequence (E+R) experiments leverage next-generation sequencing technology to track the allele frequency dynamics of populations as they evolve. While previous work has shown that adaptive alleles can be detected by comparing frequency trajectories from many replicate populations, this power comes at the expense of high-coverage (>100x) sequencing of many pooled samples, which can be cost-prohibitive. Here, we show that accurate estimates of allele frequencies can be achieved with very shallow sequencing depths (<5x) via inference of known founder haplotypes in small genomic windows. This technique can be used to efficiently estimate frequencies for any number of bi-allelic SNPs in populations of any model organism founded with sequenced homozygous strains. Using both experimentally-pooled and simulated samples of *Drosophila melanogaster*, we show that haplotype inference can improve allele frequency accuracy by orders of magnitude for up to 50 generations of recombination, and is robust to moderate levels of missing data, as well as different selection regimes. Finally, we show that a simple linear model generated from these simulations can predict the accuracy of haplotype-derived allele frequencies in other model organisms and experimental designs. To make these results broadly accessible for use in E+R experiments, we introduce HAF-pipe, an open-source software tool for calculating haplotype-derived allele frequencies from raw sequencing data. Ultimately, by reducing sequencing costs without sacrificing accuracy, our method facilitates E+R designs with higher replication and resolution, and thereby, increased power to detect adaptive alleles.

## Introduction

A major barrier to understanding the genetic basis of rapid adaptation has been the lack of robust experimental frameworks for assaying allele frequency dynamics. Recently, evolve and re-sequence (E+R) experiments^1^, which leverage next-generation sequencing technology to track real-time genome-wide allele frequency changes during evolution, have become a powerful step forward in studying adaptation^2^. In most E+R studies, replicate populations are evolved over tens to hundreds of generations in an artificial or natural selection regime and allele frequency measurements from multiple time-points are used to identify genomic targets of selection. To date, E+R approaches have already been successfully applied in a variety of model systems, including RNA molecules, viruses, *Escherichia coli, Saccharomyces cerevisiae, C. elegans* and *Drosophila melanogaster*^*3–8*^. The ability to concurrently observe both phenotypic and genomic changes across multiple systems offers the potential to answer long-standing questions in molecular evolution. Careful analysis of the patterns and magnitude of allele frequency change may reveal the extent of the genome that is under selection, how interacting alleles contribute to adaptive traits, and the speed of adaptation in different evolutionary regimes.

Crucially, however, the power to address such questions depends on the replication, time-resolution, and accuracy of allele frequency trajectories, with tradeoffs between these often incurred due to high sequencing costs. Recommended E+R schemes with even minimal power to detect selection involve sampling tens to hundreds of individuals from at least three replicate populations over a minimum of ten generations^9,10^. Since individual-based, genome-wide DNA sequencing at sufficient coverages is generally cost-prohibitive, most E+R studies rely instead on pooled sequencing^11–14^ of all individuals sampled from a given time-point and replicate. While this approach sacrifices information about individual genotypes and linkage, pooled sequencing has been shown to provide a reliable measure of population-level allele frequencies^15,16^. Still, forward-in-time simulations suggest that each pooled sample must be sequenced at a minimum of 50x coverage to detect strong selection (s > 0.1) and even higher coverage to detect weaker selection (s = 0.025)^10^. Given that optimized experimental designs often involve >100 samples, total costs for *D. melanogaster* E+R experiments that achieve reasonable detection power can reach well above $25,000 at current sequencing costs. Thus, achieving sufficient accuracy remains a major limiting factor in capitalizing on the promise of E+R.

The short timescales for which E+R is most appropriate may, however, facilitate ways to reduce sequencing costs without sacrificing experimental power. First, there is a growing body of evidence that in sexual populations, the bulk of short-term adaptation, especially in fairly small populations, relies on standing genetic variation rather than new mutations^4,13^. Many E+R schemes involve experimental populations derived from a fixed number of inbred founder lines^6,17,18^, so the identity, starting frequency, and haplotype structure of all segregating variants are often either already well-known or can easily be obtained by sequencing each founder line. Tracking only the frequencies of these validated variants can still provide enough power to detect selection, while no longer requiring the high depths of sequencing needed to differentiate new mutations from sequencing errors.

Second, at short timescales haplotype structure can be leveraged to provide more accurate allele frequency estimates. In the time frame of most E+R experiments, recombination does not fully break apart haplotype blocks and disrupt linkage, and thus genomes in an evolving population are each expected to be a mosaic of founder haplotypes. In this scenario, recently developed haplotype inference tools^19–24^ can integrate information from sequencing reads across multiple nearby sites to efficiently infer the relative frequency of each founder haplotype within small genomic windows. These haplotype frequencies can then be used as weights to calculate pooled allele frequencies for local segregating variants. With this approach, the accuracy of an allele frequency estimate depends less on the number of mapped reads at the individual site, and instead relies on the discriminatory power of all mapped reads in the surrounding genomic window when inferring haplotype frequencies. Haplotype inference methods such as harp^23^ have been shown to accurately predict haplotype frequencies at coverages as low as 25x, and simulations of pool-seq data from a small genomic region at fixed read depths indicate that the use of haplotype frequency information increases the power to detect selection compared to raw allele frequencies alone^25^. However, these tools have not yet been used to infer allele frequencies from real pooled samples in an E+R framework, nor has a thorough analysis been performed to fully examine how the accuracy of haplotype-derived allele frequency estimates scales with empirical pooled coverage, across many parameters relevant for E+R.

Here, we focus on defining the accuracy of haplotype-derived allele frequencies (HAFs) specifically in the context of E+R experiments, in which populations are generally initiated from tens of founder lines and are evolved for tens of generations. Since haplotype inference will be affected by 1) read depths throughout genomic windows, 2) recombination events, and 3) missing founder genotypes, we begin by leveraging both simulated and experimental data to assess how the accuracy of HAFs scales with each of these parameters. To do so, we introduce a new metric, ‘effective coverage’, that equates the error from HAF estimates to the expected error from binomial sampling during pooled sequencing at a given read depth. We find that haplotype inference can significantly increase the accuracy of allele frequency estimates over multiple generations of recombination with selection and with varying completeness of founder genotype knowledge. Finally, we extend these simulations in *D.melanogaster* to accurately predict effective coverage in other model organisms, such as *C.elegans*. We show as a proof of principle that a simple linear model can predict effective coverage with an R^2^ value of 0.875 and only requires four parameters: generations of recombination between population founding and sampling, average recombination rate, percent of unknown founder genotypes, and empirical read depth of the sequenced sample. Additionally, to facilitate the use of haplotype inference in E+R experiments, we introduce a software tool to calculate HAFs, HAF-pipe, (https://github.com/petrov-lab/HAFpipe-line) and a shiny app for predicting HAF accuracy in any model organism (https://ec-calculator.shinyapps.io/shinyapp/). We conclude our findings by offering recommendations about the most powerful way to integrate haplotype inference into E+R experimental schemes.

## Results

### Overview of HAF calculation method

Using haplotype inference for E+R experiments requires a genotyped founder set of inbred lines. Here, we begin by focusing on populations derived from a founder set of 99 sequenced *D. melanogaster* inbred lines (see Methods, ‘Establishing and sequencing founder set’), and for simplicity limit our analysis to the 283,437 known segregating biallelic sites on chromosome 2L. In the analyses below, we estimate raw and haplotype-derived allele frequencies (referred to as raw AFs and HAFs, respectively) for real and simulated pools of ∼100 individuals sampled from evolved populations derived from these founder haplotypes.

Each sample is first subjected to pooled sequencing and all reads are mapped to the *D. melanogaster* reference genome. Raw AFs at each of the 283k sites are simply calculated as the fraction of mapped reads containing the alternate allele, after removing reads with neither the reference nor the alternate allele. To calculate HAFs, haplotype frequency estimation is performed with harp^23^, a haplotype inference tool that uses sequence identity and base quality scores from each sequenced read in a bam file, as well as a table of founder genotypes, to obtain maximum likelihood estimates of haplotype frequencies in discrete chromosomal windows. We determined that missing calls in the founder genotype table can bias haplotype frequency estimation, and therefore, we first impute all missing genotypes before running harp (Supplemental Text, Supp. Fig. 1). The frequency of each founder haplotype is inferred within sliding genomic windows (Figure 1, Methods) with extensive overlap to mitigate erroneous haplotype frequency assignments at the edges of inference windows. After inferring founder haplotype frequencies, we calculate HAFs at each SNP site by taking the weighted sum of local haplotype frequencies for founders containing the alternate allele (Figure 1, Methods).

**Figure 1.**
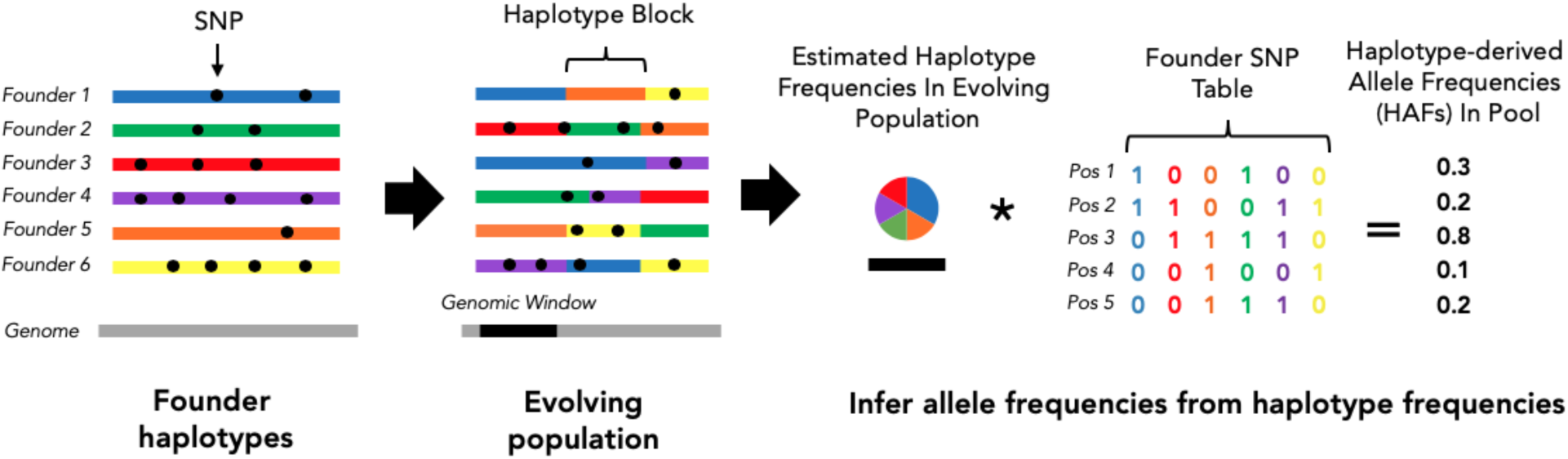
Overview of HAF calculation method. In an evolve-and-resequence experiment, the evolving population at any time point can be considered a mosaic of founder genomes. If founder genotypes are known, the relative frequency of each founder haplotype can be inferred within small genomic windows by leveraging pooled sequencing data and existing bioinformatic tools (i.e. harp). Allele frequencies can then be calculated from the weighted sum of founder haplotypes, rather than directly from mapped reads at each site.

To determine the accuracy of HAFs and raw AFs, estimated allele frequencies were compared to ‘true’ allele frequencies derived from the known composition of founder haplotypes in the sample. Chromosome-wide accuracy of HAFs and raw AFs was quantified using a new metric, effective coverage, which represents the theoretical coverage at which the root-mean-square error (RMSE) from binomial read sampling equals the RMSE from observed allele frequency estimates (Supp. Fig. 2). While expected error from extreme allele frequencies is lower than that of intermediate allele frequencies under a binomial model, by taking the ratio of the expected error and the observed error for the same set of true allele frequencies, effective coverage values do not depend on the underlying allele frequency spectrum (Methods). Note that while this metric specifically focuses on variance from the sampling of reads from pooled chromosomes, in practice, the ability of both HAFs and raw AFs to accurately reflect true population-level allele frequencies will also depend on variance from the sampling of individuals from the population. This independent source of error has however been well-treated elsewhere^26,27^ and will not be impacted by haplotype inference.

In the following sections, we explore how the accuracy of HAFs differs from raw AFs, and how it scales with empirical coverage, number of generations of recombination, and missing founder genotypes.

**Figure 2.**
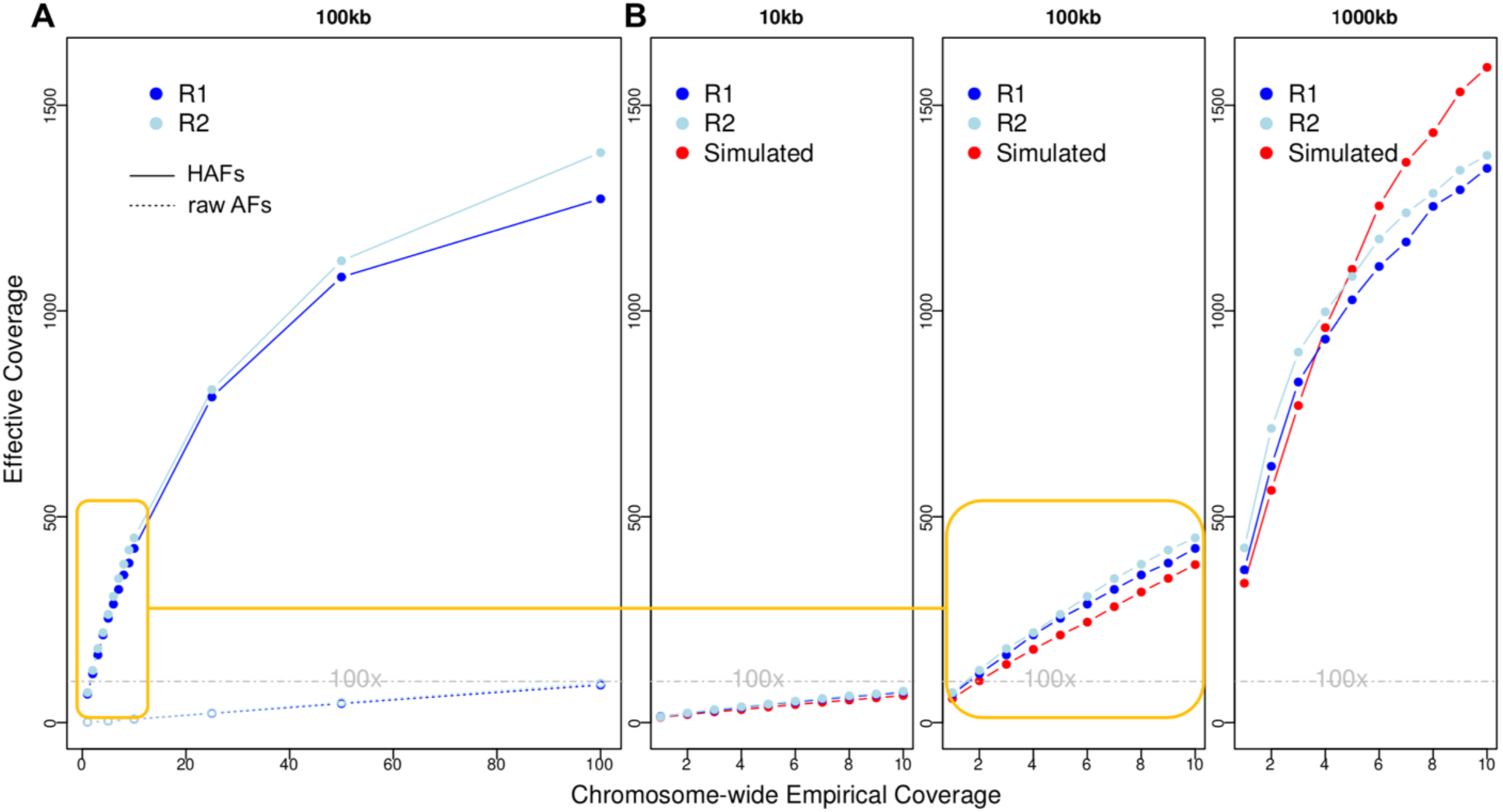
Accuracy of HAFs and raw AFs for biological and simulated samples**. A)** Effective coverage of allele frequencies estimated with and without haplotype inference for the two biological replicates down-sampled to empirical coverages from 1-100x. (R1=replicate 1, R2=replicate 2; HAFs calculated with 100kb inference windows) **B)** Effective coverages of HAFs for biological replicates (blue) and simulated samples (red) using 10kb, 100kb, or 1000kb inference windows at empirical coverages of 1-10x. Orange line indicates a zoomed-in section of the same biological replicate values as shown in A. In all panels, a dashed line of 100x indicates the accuracy threshold required to detect strong selection^9^.

### Haplotype inference significantly increases the accuracy of allele frequency estimations

We begin by analyzing the very simplest scenario, in which a sample consists of un-recombined chromosomes, one from each founder line. To examine the accuracy of HAFs and AFs in this scenario, we created two biological replicate samples of 99 *D. melanogaster* individuals, each from a different homozygous inbred strain (Methods) and performed high-coverage pooled sequencing of each sample. True allele frequencies for each sample were then calculated, incorporating estimates of uneven pooling during sequencing (Supp. Fig. 3, Supp. Text).

The accuracy of HAFs depends on the power to estimate haplotype frequencies, which in turn is affected by the coverage of mapped reads throughout the genomic window used for haplotype inference. In order to compare the accuracy of HAFs to raw AFs and test how each scales with empirical coverage, reads from the two biological replicates originally sequenced at ∼140x were down-sampled to chromosome-wide empirical coverages of 1x to 100x, and then used to calculate the effective coverage of estimated allele frequencies for each replicate (Figure 2). Haplotype inference was initially performed using 100kb sliding windows, and accuracy was assessed only at the 27k sites with known genotype information for every founder. As expected, effective coverage of both HAFs and raw AFs is similar within the two biological replicates, and increases with higher chromosome-wide empirical coverage. Yet for all empirical coverages tested, HAFs have strikingly higher effective coverages than raw AFs. This substantial gain in accuracy from haplotype inference was most prominent at lower empirical coverages, with a >40-fold increase in accuracy at 10x empirical coverage (from 10x to effectively >400x). At higher empirical coverages, haplotype inference appears to produce diminishing returns and effective coverage begins to plateau.

We next tested the effect of using smaller (10kb) or larger (1000kb) windows for haplotype inference at empirical coverages up to 10x. We find that as expected, larger window sizes produce the most accurate HAFs, since more reads are available to infer haplotype frequencies in each window. Specifically, 1000kb HAFs derived from empirical coverages of 1x and 5x reached effective coverages of >400x and >900x, respectively.

We also confirmed that similar results would be achieved by simulated samples with known sources of error. To do so, we simulated pooled synthetic reads with a standard Illumina sequencing error rate of 0.002^28^ and corresponding base quality scores^23^ from the same proportions of the 99 founder lines included in both biological replicates, and calculated effective coverage with the same empirical coverages and window sizes as above. Effective coverages for these simulated samples closely mirror effective coverages obtained from matched experimental samples (Figure 2A-B). Slight differences at higher empirical coverages and larger window sizes are most likely caused by compounded experimental error from DNA extractions, PCR reactions and sequencing, as well as ambiguity in the ‘true’ genotypes estimated for individually sequenced lines. We explore the effects of founder genotype ambiguity further below (see “HAF accuracy is impacted by missing founder genotypes”).

Together, these results suggest that HAFs derived from multiple biological samples sequenced at low empirical coverages can be orders of magnitude more accurate than raw AFs, and that simulated samples capture the magnitude of this effect quite well. In the following analyses we focus on simulated data from these 99 founder lines in order to precisely and reliably test how recombination and missing founder genotypes affect HAFs in realistic E+R scenarios.

### Incorporating recombination and selection over short time scales

In the first section we showed that HAFs are accurate for samples of unrecombined chromosomes with very similar allele frequencies to the founder population. However, in a realistic E+R scenario sampled chromosomes will be recombined mosaics of the founders and selection may substantially shift allele frequencies over time. Thus, in the remainder of our analyses, we incorporate selection and recombination using a forward-in-time simulator, forqs^29^. We simulated recombination for 50 generations in a population of 1,000 randomly-mating individuals using a *D. melanogaster* recombination map^30^, and tracked the breakpoints and haplotype of origin for all recombined segments at every generation. At specific generations, we randomly selected 200 recombined chromosomes (i.e. 100 diploid individuals) from the simulated population and reconstructed the full sequences of these ‘sampled’ chromosomes from corresponding segments of the 99 individually sequenced founder haplotypes. Reads were simulated from the pooled set of 200 reconstructed chromosomes.

We tested the accuracy of HAFs in three different selection regimes: 5 randomly chosen sites with selection strength s=0.025 (i.e. weak selection), 10 randomly chosen sites with selection strength s=0.025, and 5 randomly chosen sites with selection strength s=0.1 (i.e. strong selection). In each case, the selected sites contributed additively to a single quantitative trait (Methods). We ran each simulation in three rounds, picking a different set of selected sites in each round, and simulated 5 replicate populations for each round. We calculated HAFs for each simulated sample, adjusting the window size for haplotype inference each generation based on the expected length of unrecombined haplotype blocks (Methods, Supp. Fig. 4). As recombination proceeds, these windows become smaller, with fewer reads available to inform haplotype frequency estimation, and thus we expect that accuracy will decline.

Our results show that while accuracy does decline over time, HAFs in general maintain >100x effective coverage even after 50 generations in the presence of 5-10 selected sites per chromosome (Figure 3A). We note that the three selection regimes tested all perform comparably well, though effective coverage is slightly higher with more selected sites (i.e. 10 vs. 5) and larger selection coefficients (i.e. s=0.1 vs. s=0.025). To assess the utility of HAFs for longer-term experiments, we also conducted three separate simulations with weak selection and 5 selected sites for 200 generations, and note that effective coverage of HAFs remains above the 100x threshold for detecting strong selection across 100-150 generations of recombination (Supp. Fig. 5). Overall, we find that increasing empirical coverage has a diminishing returns effect on accuracy, while decreasing the generations of recombination has an approximately linear effect on accuracy.

**Figure 3.**
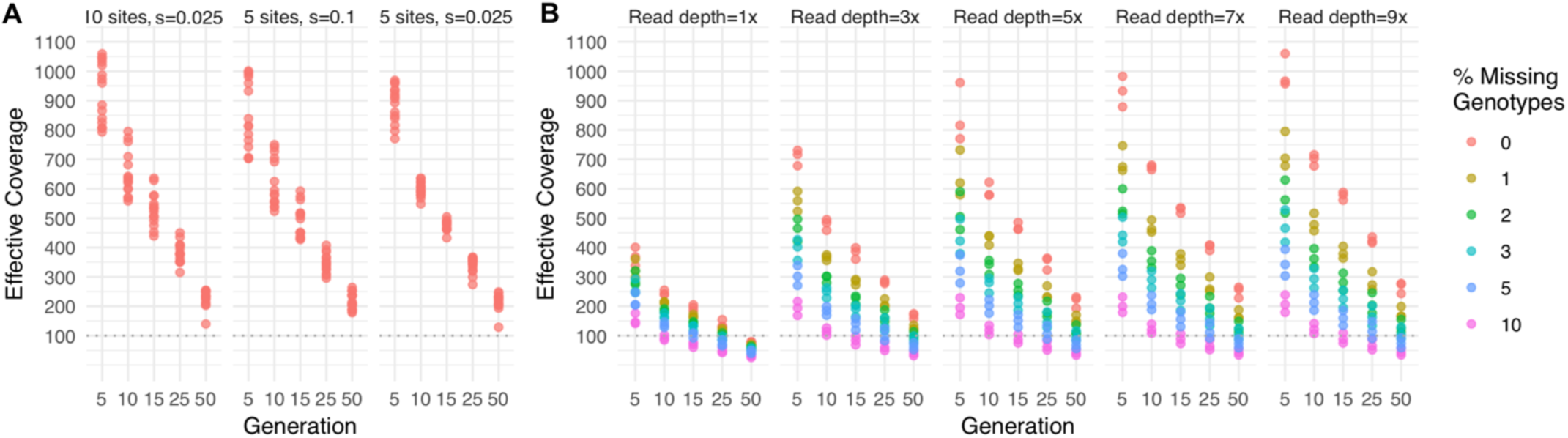
Effective coverage for recombination simulations with 99 inbred founder lines. In each simulation, recombination was simulated for a population of 1,000 individuals initiated from a panel of 99 fully sequenced inbred *D. melanogaster* lines, with a randomly chosen set of selected sites from among the 283,437 segregating sites on chromosome 2L. At multiple generations, 100 recombined individuals were sampled *in silico* from each population for simulated sequencing, HAF calculation, and error estimation. **A**) 5 or 10 sites were under weak or strong selection (panels), all sequencing was simulated at 5x, and HAFs were calculated with no missing genotypes. **B**) 5 sites were under weak selection, sequencing was simulated at multiple read depths (panels), and HAFs were calculated with various fractions of missing founder genotypes (color).

### HAF accuracy is impacted by missing founder genotypes

In addition to the effects of recombination and selection on HAF accuracy, ambiguity in the genotypes of founders (either due to missing genotype information or residual heterozygosity) can also present challenges. We tested how setting founder genotypes to be ambiguous (i.e. from a called ‘A’ allele to an uncalled ‘N’) influences the accuracy of HAFs. For each simulation, 1-10% of all genotype calls in the founder table were randomly selected to be assigned as ‘N’. Genotypes at these sites were then imputed prior to estimating haplotype frequencies (Supplemental Text) and calculating HAFs. For each percentage, three rounds of simulation were performed.

We find that missing genotype calls can significantly reduce effective coverage (Figure 3B). However, the vast majority of the parameter space tested — 73.3% of all simulations — still yielded effective coverage values greater than 100x. Furthermore, 92.1% of simulations with less than 10% missing genotypes and fewer than 50 generations of recombination yielded effective coverage values greater than 100x.

Importantly, these high overall chromosome-wide effective coverages extend to the selected sites themselves. Even with moderate levels of missing founder calls (up to 3 % of missing sites), HAFs at selected sites still track closely with true allele frequencies; this is crucial for correctly detecting alleles under selection in E+R (Supp. Fig. 6).

### Estimating effective coverage with different founders sets and other model organisms

Finally, we explored how the utility of HAFs may extend to other founder sets with known SNPs and known recombination rates. Specifically, we tested whether a simple linear model based on the simulations above could accurately predict effective coverage for other experimental designs and other model organisms. The regimes tested in the simulations above can be collapsed into two independent parameters that affect HAF accuracy: 1) the number of reads used for haplotype inference in each window and 2) the percent of founder genotypes that are missing. The first parameter is a combination of SNP density, read depth, and window size – while the window size itself is a function of the recombination rate and the number of generations of recombination since population initiation.

We calculated regression coefficients of a log-linear model (Supp. Fig. 7) using all simulations described in the sections above, which focused on a single founder set across many experimental regimes. To test whether this model could accurately predict HAF accuracy for other founder sets, we simulated three rounds of evolutionary trajectories (with 5 weakly selected sites in each round) for two entirely different founder sets composed of 1) a widely-used reference panel of 205 *D. melanogaster* lines known as the DGRP^31^, and 2) 100 genotyped *C. elegans* strains from a reference panel known as CeNDR^32^. For each founder set we simulated pooled samples for various generations and empirical coverages and calculated HAFs with various levels of missing founder information. We then used our model (trained only on the original simulations) to predict effective coverage for each of these new samples (Figure 4A).

**Figure 4.**
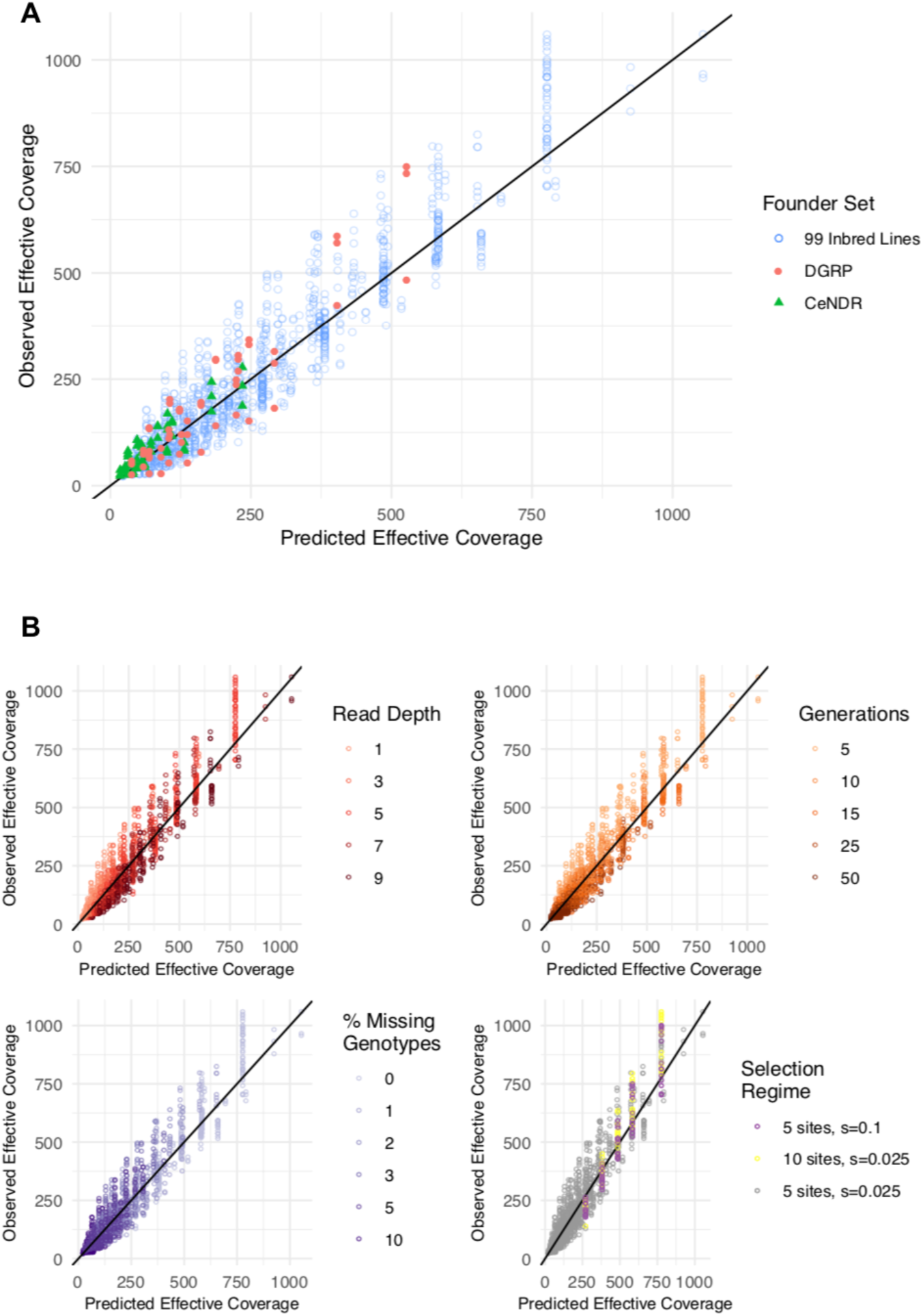
Observed effective coverage vs. effective coverage predicted by a simple 2-parameter log-linear model. **A**) A model built on samples from the simulated experimental evolution of 99 inbred *Drosophila melanogaster* lines described in the sections above was used to predict effective coverage in the simulated experimental evolution of different founder panels (205 DGRP lines) and different founder species (100 CeNDR lines). **B**) Effective coverage was well-predicted across a range of simulation parameters, including read depth, number of generations, % missing founder genotypes, and selection regimes.

We find that across all samples tested, our simple 2-parameter model has an R^2^ value of 0.875. Predicted effective coverage values for simulations with the original 99-lines founder set differ from true effective coverage values by 25% on average, with the largest source of error due to random effects between simulation rounds. Overall accuracy for DGRP and CeNDR samples was slightly lower, with average deviations of 37% in both founder sets. We confirmed that the model does not systematically over- or under-predict the effective coverage of DGRP samples or CeNDR samples nor any other set of parameters included in our simulations (Figure 4B). This suggests this model is broadly applicable for predicting HAF accuracy across many founder sets. Thus, given a set sequencing budget and a founder set with known sequencing quality, this model may be useful as a guideline for devising experimental schemes and distribution of resources that would maximize detection power. To this end, we have created a shiny app to help experimentalists predict effective coverage for their particular set of parameters (freely available at https://ec-calculator.shinyapps.io/shinyapp/), as well as a table of requirements for HAF estimation (Figure 5).

**Figure 5.**
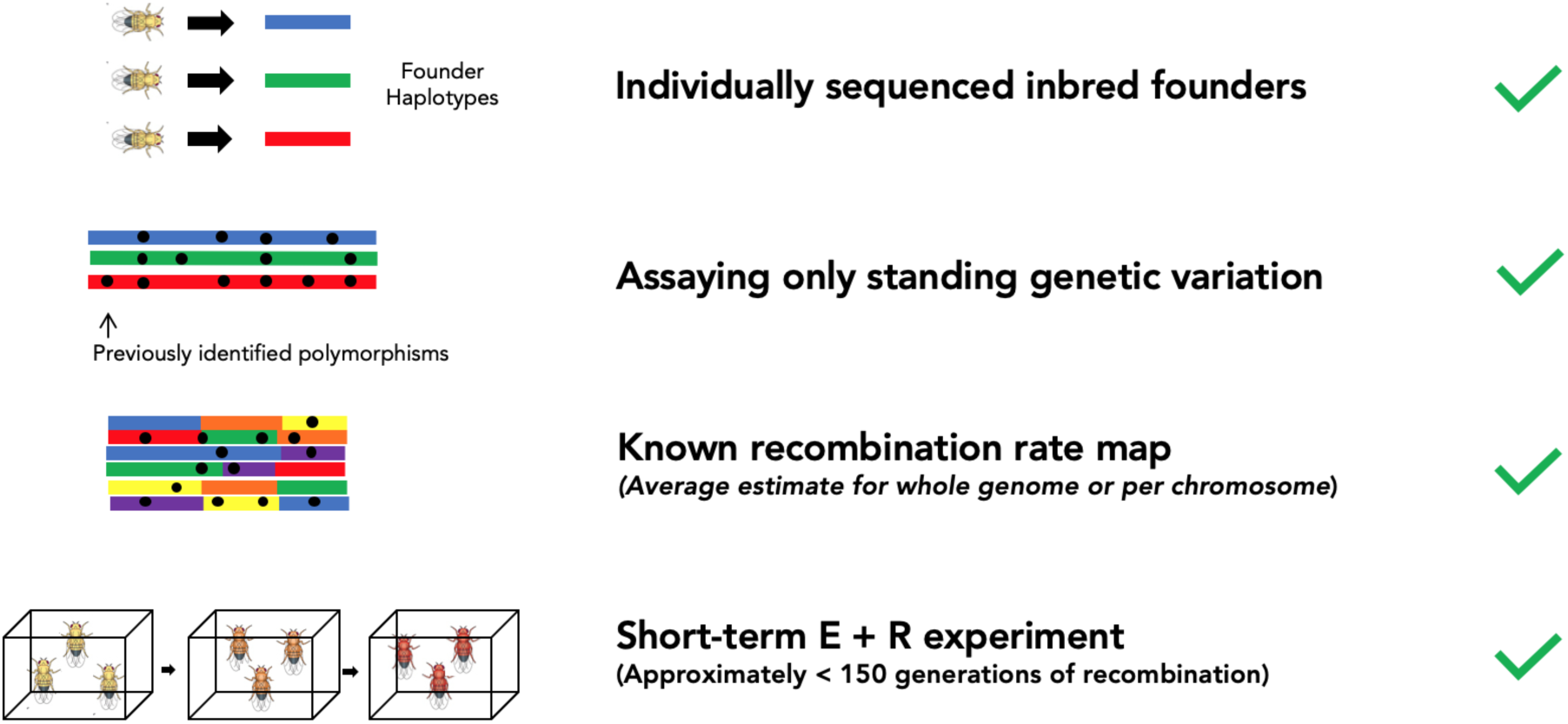
Schematic of requirements for using HAFs to estimate allele frequencies in E+R experiments. Recommendations of timescale are based on simulations with *D. melanogaster*

## Discussion

E+R experiments have become a powerful tool to assay the underpinnings of rapid adaptation by tracking allele frequency trajectories within populations over time. Previous studies have shown that the greatest power to detect adaptive variants comes from an optimized experimental design that tracks allele frequencies in multiple replicate populations, samples each replicate population at multiple timepoints, and maximizes the coverage of each pooled sample. Incorporating all of these factors into an E+R framework, however, can present significant financial challenges. Here, we offer a way to mitigate these high sequencing costs without sacrificing statistical power.

Our framework uses haplotype inference to increase the accuracy of pooled allele frequency estimates at low coverages. Since the accuracy of haplotype-derived allele frequencies relies on the total informative value of reads across a genomic window, rather than coverage at a single site, this approach allows us to sequence less but still maintain high accuracy in allele frequency estimations. In this vein, the window-based techniques used by HAFs have an advantage over raw AFs in that the accuracy of an individual SNP with low read depth will benefit from reads at surrounding sites in the window. Overall, our method is capable of achieving the same accuracy as would be expected from sequencing each sample at 100x (as recommended in order to reliably detect strong selection), while only requiring empirical coverage of 1x or less, bringing total sequencing costs from >$25,000 down to less than $200.

There are, however, limitations to this approach. First, as presented, this framework requires the founder population to be derived entirely from fully-inbred lines. As a result, the population dynamics of loci under selection may differ slightly from trajectories in natural populations due to the genetic diversity lost in the inbreeding process (i.e. natural haplotypes, homozygous lethal mutations, and rare variants), as well as higher levels of linkage disequilibrium compared to non-inbred lines. Reconstituting an outbred population using inbred lines, however, can be an effective way to mitigate the effects of the inbreeding process, and has been experimentally shown to have negligible bias and effect on adaptive dynamics^33^. Alternatively, the use of inbred lines may not be necessary with more sophisticated founder sequencing approaches that incorporate phasing. Newer long-read technologies may make this an achievable reality for a number of systems in the near future.

Second, this approach requires a reliable and comprehensive account of the variants present in each founder line. Since previous studies recommend upwards of 100 founders, sequencing each individual founder line to a sufficiently high depth (in our work, we have found sequencing coverages >10x to be sufficient) may present a high upfront cost. However, this cost represents a one-time investment, which can be applied toward all future experiments using the same set of founders. Furthermore, a number of consortiums already maintain publically available stocks of large numbers of *Drosophila* lines and other model organisms with full, high-quality genome sequences ^31,32,34^. We anticipate that these resources will continue to rapidly expand, facilitating experiments with even greater haplotype diversity at minimal costs.

In addition, this approach is limited to studying short-term adaptation on the scale of tens of generations. In fact, an assumption of our method is that within an inference window, recombination breakpoints minimally affect the ability to accurately call haplotype frequencies. For a given window size however, this assumption becomes less valid as recombination proceeds and haplotypes blocks decay. Conversely however, decreasing the window size reduces the information used for haplotype inference, which at the extreme renders HAFs no more accurate than raw AFs. In our pipeline, we attempt to balance these effects by scaling window sizes at any generation by the expected average unrecombined fragment length. While our results here demonstrate that even with this scaling procedure, recombination will limit the ability to detect adaptation on timescales of more than tens of generations, the short-term adaptive dynamics that best fit E+R studies fall well within this range. Furthermore, it is at these short timescales, when large numbers of replicate populations are critical to reliably detect selection, that the cost savings associated with haplotype inference methods will be most beneficial.

Finally, this approach relies on tracking the trajectories of known bi-allelic polymorphisms derived from the founder population, and thus, de novo mutations will not be assayed in this framework. Nonetheless our approach should sufficiently capture the salient features of short-term adaptive dynamics, as there is a growing body of experimental evidence suggesting that selection acts primarily on standing genetic variation in sexual organisms, and that de novo beneficial mutations do not play a large role in rapid adaptation ^4,35–37^. Additionally, by tracking only known well-validated polymorphisms, the approach is largely robust to error from small non-SNP chromosomal variants such as indels.

Despite the above limitations, collectively our results show that integrating haplotype inference into future E+R experiments is a cost-effective way to achieve accuracy in allele frequency estimates, which will directly improve the ability to detect genome-wide signatures of adaptation. Consequently, we offer specific recommendations for future E+R experimental schemes that take advantage of this approach. First, each founder line should be initially sequenced to a sufficient depth that minimizes any missing genotypes. If missing genotype calls do exist in founder lines, imputing sites prior to haplotype inference can mitigate some of this error.

Together, these guidelines and the analysis above form a framework for achieving effective coverages of close to 100x with empirical coverages as low as 1x even after 50 generations of recombination in *Drosophila melanogaster*, reducing sequencing costs by 100-fold. Ultimately, these cost savings, which can be extended to experiments with a variety of model organisms, will facilitate E+R frameworks that can incorporate large numbers of replicate populations. These improvements may be crucial to the statistical power to distinguish between beneficial and neutral alleles ^38,39^ and ultimately the future of E+R as a practical and reliable experimental tool.

## Methods

### Establishment and sequencing of founder set

207 iso-female *Drosophila melanogaster* lines were established from wild individuals sampled from Maine and Pennsylvania^40^, and inbred for ∼20 generations of full-sibling mating to produce viable, fertile inbred lines. 30-50 individuals from each line were pooled for DNA extraction. Whole flies were homogenized with lysis buffer and 1mm beads, and DNA was precipitated from the homogenate before resuspension in TE buffer. Libraries were prepared with a modified Nextera protocol^41^. All samples were indexed with Illumina’s TruSeq Dual Index Sequencing Primer Kit (PE-121-1003) and pooled equimolarly into 3 sets of ∼70 samples each. Each set of pooled DNA libraries were purified using Ampure XP and size-selected to 450-500 bp with a SizeSelect E-Gel. After an additional 5 rounds of PCR, DNA libraries were purified using Ampure XP beads, quantified, and diluted to the appropriate concentration before sequencing on the HiSeq 3000. All sequences were deposited in SRA (BioProject PRJNA383555). Adapter sequences were trimmed (Trimmomatic v0.33) and overlapping reads were merged (PEAR v0.9.6), then reads were mapped (bwa v 0.7.9) to the *D.melanogaster* reference genome (v5.39) using default parameters. PCR duplicates were removed using PicardTools (v1.12). Base quality score recalibration, indel realignment, and novel SNP discovery were carried out using GATK’s HaplotypeCaller. Only bi-allelic SNPs segregating in the 99 lines pooled for resequencing in this study were used to generate a founder SNP table, simulate reads, and estimate haplotype frequencies.

### Generating Experimentally Pooled Samples

One male each was selected from each of 99 inbred strains, and all 99 individuals were pooled for re-sequencing. A second biological replicate was constructed from 99 additional individuals. DNA isolation was performed as described above. Three separate libraries were prepared from each of the two biological replicates using different library prep methods: [1] according to protocols described in Nextera DNA Library Prep Reference Guide (15027987 v01); [2] a modified Nextera protocol (as described above) and [3] a Covaris shearing protocol. Final results from the 3 library prep methods were similar. All libraries were size-selected and PCR amplified using two replicate PCR reactions and a high volume of template DNA to prevent PCR-jackpotting. DNA was purified, quantified, and diluted before sequencing on the HiSeq 3000. Raw, 150bp pair-end reads were trimmed for adapter sequences with Skewer (version 0.1.127). Read merging, mapping, and PCR duplicate removal was performed as above.

### Generating Simulated Pooled Samples

150-bp paired end pre-aligned reads were simulated from a table of founder genotypes and the *D. melanogaster* reference genome with simreads, a software tool included with the harp package^23^. All reads were simulated with an error rate of 0.2%^28^, with simulated sequencing errors receiving a lower simulated base quality score. No read trimming or PCR duplicate removal was done. All SNP tables with missing genotypes were imputed before read simulation.

### Haplotype Frequency Estimation

All haplotype frequencies were estimated with harp - Haplotype Analysis of Reads in Pools ^23^ in a two-step process in which 1) a likelihood model is built by probabilistically assigning all reads to haplotypes, and 2) maximum likelihood estimates of haplotype frequencies are calculated in discrete chromosomal windows, given local read assignments. An assumption of this method is that there are no recombination breakpoints within a window used for haplotype frequency estimation. However, with a fixed window size, this assumption breaks down as the lengths of unrecombined fragments decrease. The distribution of fragment lengths at a given generation can be modeled with an exponential distribution with rate, *λ*, equal to,

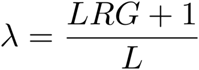

where R is recombination rate, L is chromosome length, and G is the number of generations of recombination between the initiation of the founding population and sampling. The *q*th quantile of this distribution can be calculated in R with the function qexp(*q,λ*).

We allowed window sizes to shrink over successive generations of recombination, such that only 18% of sampled unrecombined fragment lengths were expected to be smaller than the window length. Various quantiles from 5-25 were tested before choosing this parameter (see Supp Fig. 4). Note that haplotype frequencies for fully unrecombined chromosomes (Fig. 2) were evaluated in 1000kb, 100kb and 10kb windows. To further reduce error, we used overlapping inference windows, with a step size equal to 10% of the window size. Thus, the vast majority of sites fall within 10 separate overlapping inference windows. Finally, in order to balance local relevance with maximal information, we always created likelihood models in windows 10x the size of frequency estimation windows, with a step size equal to half the likelihood window size.

For reference, inferring haplotype frequencies for 99 founder lines at 283k segregating sites on chromosome 2L in 1000kb windows took 8 minutes and required 450Mb RAM for samples sequenced at 5x empirical coverage and took 15 minutes and required 860Mb RAM for samples sequenced at 10x. Using 100kb windows took 9.5 minutes / 70Mb and 17.5 minutes / 132Mb for 5x and 10x samples, respectively.

### HAF Estimations

The haplotype-derived allele frequency (HAF) for a given biallelic site was calculated by summing founder haplotypes containing the alternate allele, each weighted by their average estimated haplotype frequency in all haplotype inference windows overlapping the site. Founder haplotypes with missing genotypes were given a fractional alternate allele count equal to the mean of genotyped founders with alternate alleles.

### Accuracy Estimations Using Effective Coverage

Effective coverage was used as a metric to assess the accuracy of all HAFs and raw AFs. For a given set of allele frequency estimates *p*_*estimated*_ at *n* sites, for which true frequency *p*_*true*_ is known, we first calculate the root mean squared error (*RMSE*_*estimated*_), where

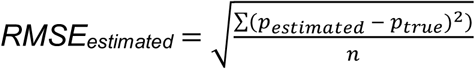

Next, we solve for the coverage *C*_*effective*_ at which *RMSE*_*theoretical*_ from binomial sampling would be equal to *RMSE*_*estimated*_, where

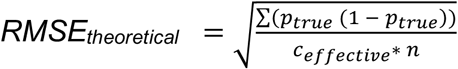

Solving for *C*_*effective*_ yields,

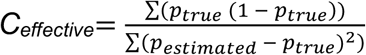

which is the theoretical coverage at which binomial sampling of reads would be expected to contain the observed amount of error from estimated frequencies.

### Recombination

Forward-in-time simulations of recombination were performed with the software tool forqs^29^ using a *D. melanogaster* recombination map^30^. forqs simulates recombination of haplotype chunks for chromosomes of user-specified lengths for a randomly mating population, using a user-supplied recombination map, and non-overlapping generations. As a conservative metric, in our simulations we referred to the female *D. meanogaster* recombination rate. Since male *D melanogaster* do not undergo recombination, our estimates of the number of recombination events per generation are higher than that expected in real populations and our estimates of effective coverage serve as a lower bound on effective coverage expected at the same number of generations in real populations. Three rounds of simulation were performed for each selection regime. In each round, an initial population of 1,000 individuals was created, with each individual assigned to a randomly selected homozygous founder strain. 5-10 sites were randomly chosen to be under selection and the genotypes of each individual (determined by the genotype of the corresponding founder strain) at these sites was supplied to forqs via an ms file. Homozygous reference, heterozygous, and homozygous alternate genotypes were assigned fitness advantages equal to 0, *s*, or 2*s* respectively, where s was a specified selection coefficient (either s=.025 or s=.1 in our simulations). The chosen loci each contributed independently to a single additive trait, with environmental variance equal to 0.01. At each generation, a fitness value was calculated by forqs for each individual based on their genotypes at the selected sites, with fitness decaying linearly with distance from the optimum trait value of 1. Individuals were selected to contribute to the next generation probabilistically based on their fitness value. Recombination breakpoints were simulated for evolutionary trajectories up to 50 generations in 5 replicate populations with a constant population size of 1,000 individuals. Within each round, each replicate contained the same selected sites and selection coefficient. At specific generations, 100 sets of recombination breakpoints (each representing a pair of evolved ‘chromosomes’) were randomly selected from the forqs output and were used to reconstruct ‘sampled chromosome genotypes’ from corresponding segments of the 99 founder genotype calls. This set of sampled genotypes was used to directly calculate ‘true’ allele frequencies for the sampled pool and was also used as input for read simulations with simreads. The resulting reads were then used for HAF calculation.

### Generating a predictive model of effective coverage

While we observed non-linear relationships between effective coverage and both parameters, the log-log relationships were fairly linear. This suggested that a reasonable simple model would have the following format:

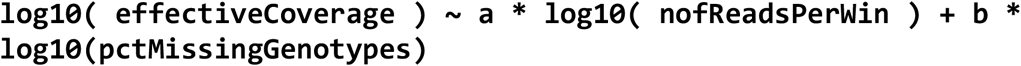

We used the R function ‘nls’ to solve for the coefficients *a* and *b* in this formula, using all *Drosophila melanogaster* simulations described in the sections above.

### HAFs with an alternate founder set

For the DGRP founder set, SNP information was obtained for 205 strains initially isolated from Raleigh, NC that were independently sequenced as part of freeze 2 of the *Drosophila* Genetic Reference Panel (DGRP)^31^. Genotype data was downloaded directly from http://dgrp2.gnets.ncsu.edu. For the *C. elegans* founder set, a soft-filtered VCF file (*v. 20170531*) of genotype calls for 249 sequenced strains ^32^ was downloaded from the CeNDR website (https://www.elegansvariation.org/data/release/20170531), and was converted to a SNP table including genotypes for 100 randomly selected lines at all segregating biallelic SNP sites.

After constructing the appropriate SNP table, read simulation, haplotype inference and effective coverage calculations were carried out as described in the sections above.

### Code Accessibility

Scripts to calculate HAFs are available at https://github.com/petrov-lab/HAFpipe-line. At minimum, the pipeline requires a) called biallelic variants from sequenced founder lines, and b) mapped reads from one or more pool-seq samples, and uses harp for haplotype inference.

### Statement on Data and Reagent Availability

Sequence data from seasonal strains is available at SRA (BioProject PRJNA383555) and genotype data is available at *https://github.com/petrov-lab/HAFpipe-line/blob/master/99.clean.SNP.HARP.segregating.gz*. Strains are available upon request. Code used to generate the simulated data is provided at https://github.com/petrov-lab/HAFpipe-line/tree/master/simulations.

## Supplemental Text

### Incorporating uneven pooling of individuals produces more realistic estimates of true allele frequencies

Our ability to measure the accuracy of HAFs and raw AFs depends on our ability to determine the true contribution of each pooled individual. Since uneven pooling is a source of error known to affect pool-seq samples^15^, we estimated the relative contribution of DNA from each individual by calculating the average genome-wide allele frequency at sites private to each founder. While each founder could be detected in the pool, we found substantial variation in their relative representation (Supp. Fig. 3). ‘True’ frequencies for the experimental pooled sample were thus calculated by weighting founders known to contain the alternate allele by their estimated representation in the pool. We assessed whether these ‘true’ allele frequencies were better recapitulated by experimental reads than ‘true’ allele frequencies calculated without incorporating uneven pooling at all fully genotyped sites (both private and common). We found that the effective coverage using unevenly pooled weighted values (126x) was higher than the effective coverage assuming evenly pooled individuals (120x). We used these same estimates of uneven pooling to simulate reads in uneven proportions from different haplotypes for the synthetic sample as well.

### Imputing missing founder genotypes increases the accuracy of HAFs

While missing information can be accommodated by many haplotype inference tools (i.e. an N in place of a missing call), it is unclear how missing calls affect inference accuracy, and what the best practices should be when missing calls are present in the reference founder set.

We first examined whether haplotype frequencies estimated for founders with many missing calls or few missing calls systematically deviated from an expected haplotype frequency of 0.101 (1/99). We found that across individual inference windows, there was a clear negative correlation between the number of missing calls per founder, and the haplotype frequencies estimated for that founder (Supp. Fig. 1). To determine whether the observed skewed haplotype frequencies were directly associated with the presence of missing sites, we tested whether imputing genotype calls for missing sites would reduce bias in haplotype frequency assignment. While a number of sophisticated methods for imputing rare SNPs do exist ^42–44^, and may in some cases improve HAF accuracy, here we used a simple approach. To perform imputation, at each site we first calculated the allele frequency among called founder genotypes and used this value as a probability for assigning genotypes to missing calls. We found that imputation significantly reduced the skewed haplotype frequency distribution by 4-6-fold for all empirical coverages and window sizes tested. We expect that imputation with more advanced tools would achieve even better results.

We next examined how imputation of haplotype frequencies can impact the overall accuracy of HAFs. We also confirmed that haplotype inference using imputed calls produced more accurate HAFs than using a subset of sites with no missing calls. Thus, we include imputation as a key step in our analysis pipeline.

## Supplemental Figures

**Supplemental Figure 1.**
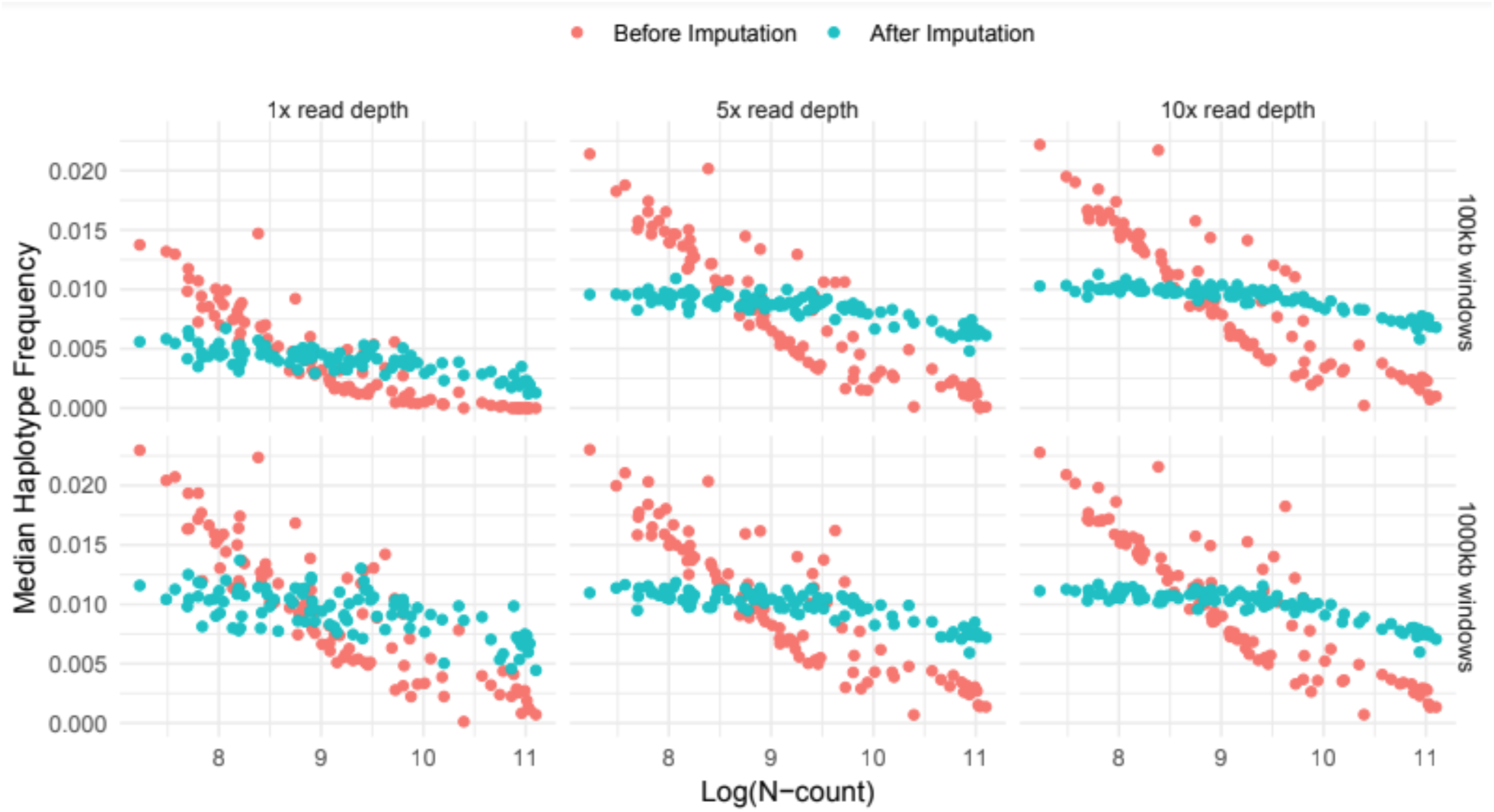
Median haplotype frequency across all windows on chromosome 2L for each founder (n=99), calculated with different window sizes and empirical coverages. Haplotype frequencies calculated before imputation (red circles) and after imputation (blue circles) are plotted as a function of the log of the total number of ambiguous genotypes (aka “N-count”). Best fit lines for each dataset were calculated with standard linear regression.

**Supplemental Figure 2.**
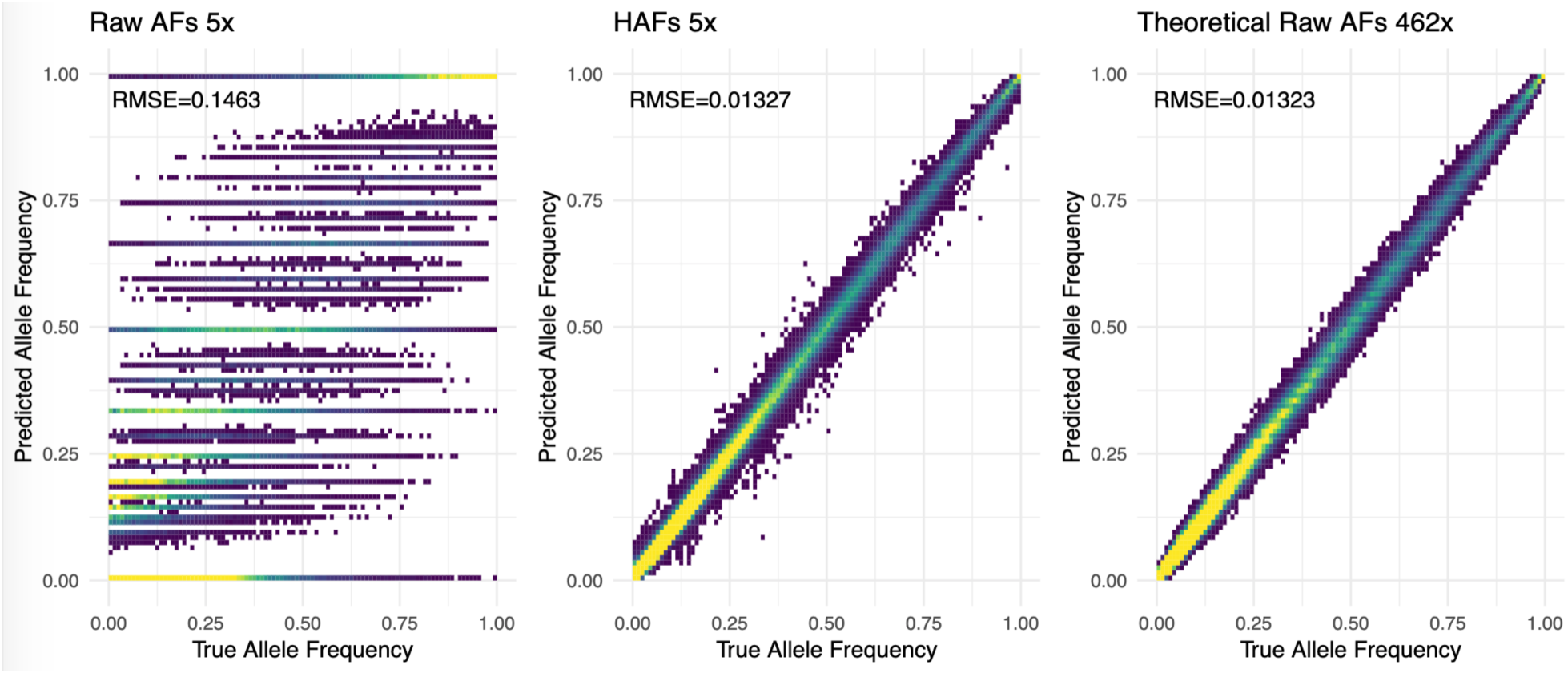
An example of true and predicted allele frequencies at each segregating site on chromosome 2L, where predicted frequencies are calculated either from A) raw mapped reads at 5x empirical coverage, B) HAFs at 5x empirical coverage, C) simulated binomial sampling of reads at 462x coverage. Color represents density of points. RMSE for each set of predictions is indicated in the top left of each panel. Note that RMSE for panels B and C are very similar; this equivalence forms the basis of assigning an ‘effective coverage’ of 462x to the estimated allele frequencies in panel B.

**Supplemental Figure 3.**
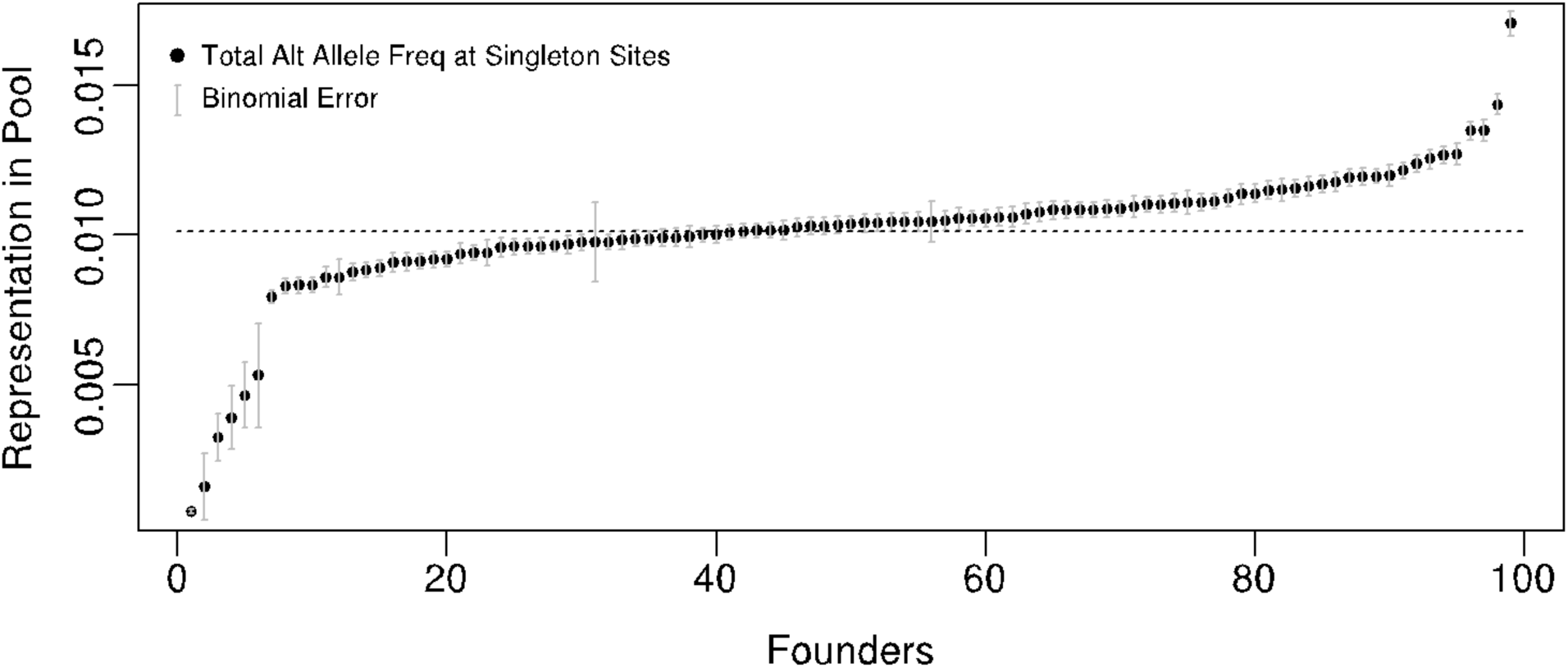
Contribution of DNA from each pooled individual in experimental replicate 1, estimated by average genome-wide allele frequency across all singleton sites. The dashed line represents theoretical expectation for evenly pooled individuals. Error bars represent total expected binomial error, given total read depth at all singleton sites for a given founder.

**Supplemental Figure 4.**
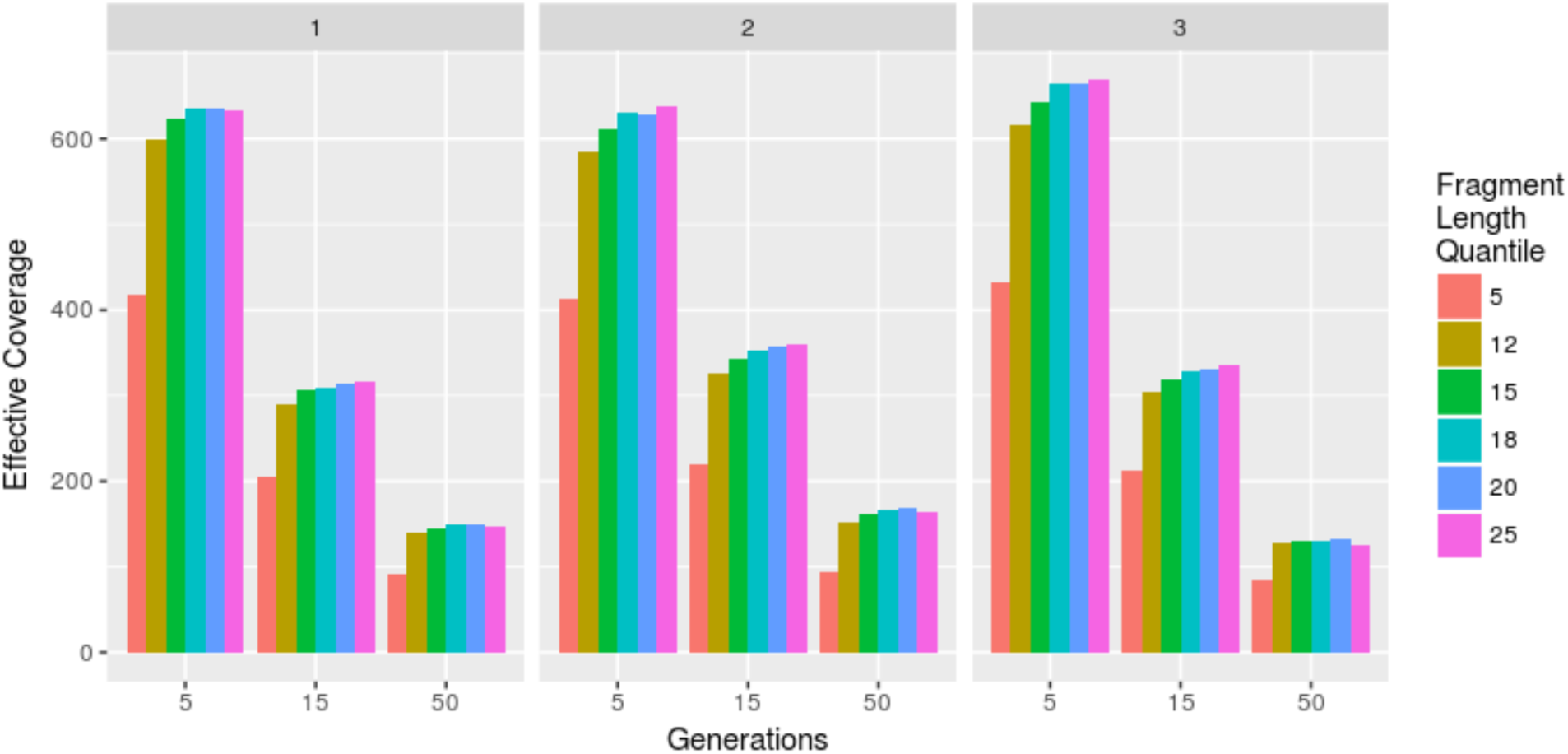
Effective coverage was calculated for samples simulated at 5x empirical coverage after 5,15, and 50 generations of weak selection, with a founder genotype table missing 1% of calls, using various window sizes for haplotype inference. Colors correspond to the quantiles of the expected exponential distribution of unrecombined fragment lengths that were used as the window size for haplotype inference. Each panel (1-3) represents results from a different simulation round, using a different set of selected sites.

**Supplemental Figure 5.**
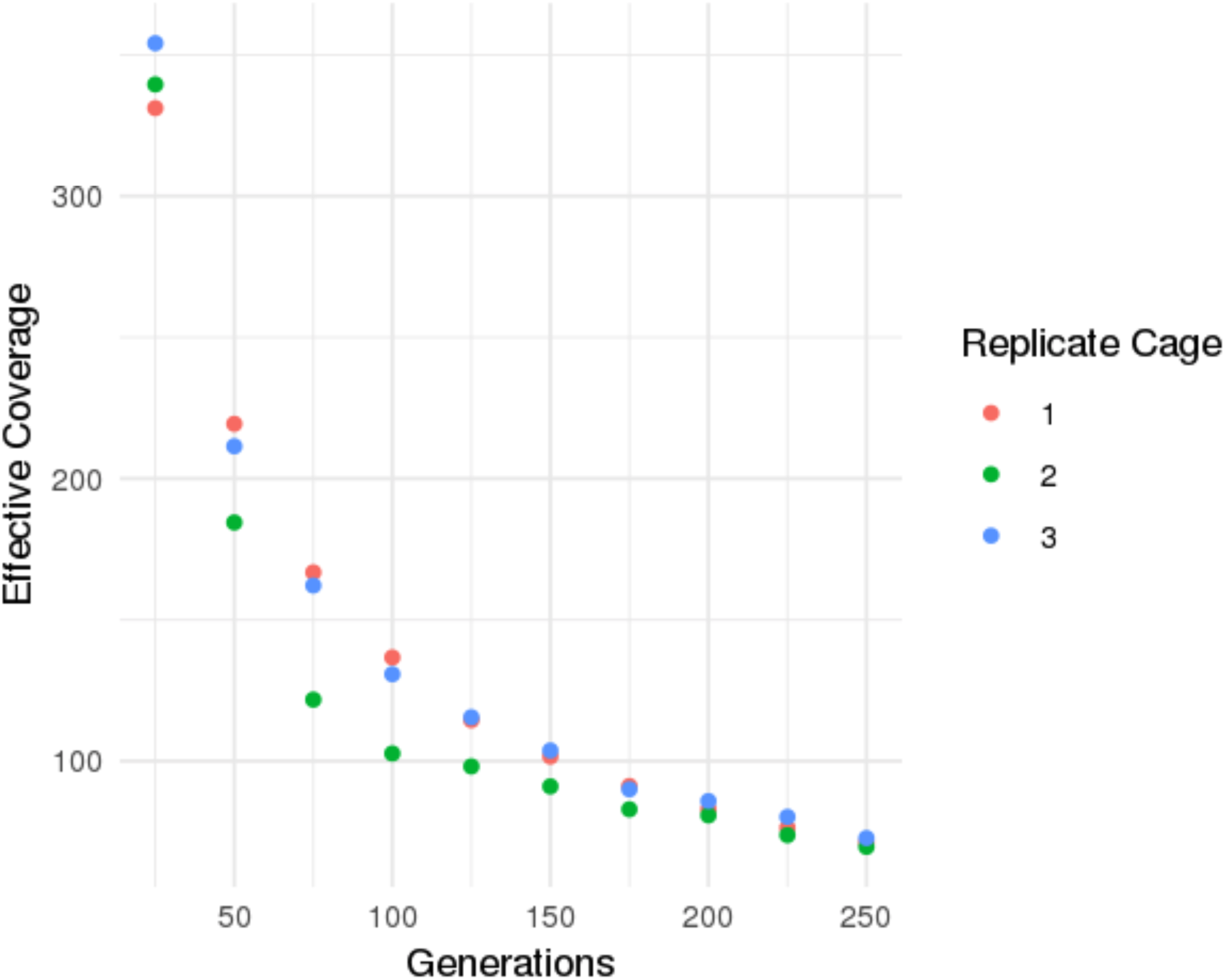
Effective coverage for 3 separate simulated long-term experiments each with 5 randomly selected sites under selection (S=0.025), simulated empirical coverage of 5x, and no missing founder genotypes.

**Supplemental Figure 6.**
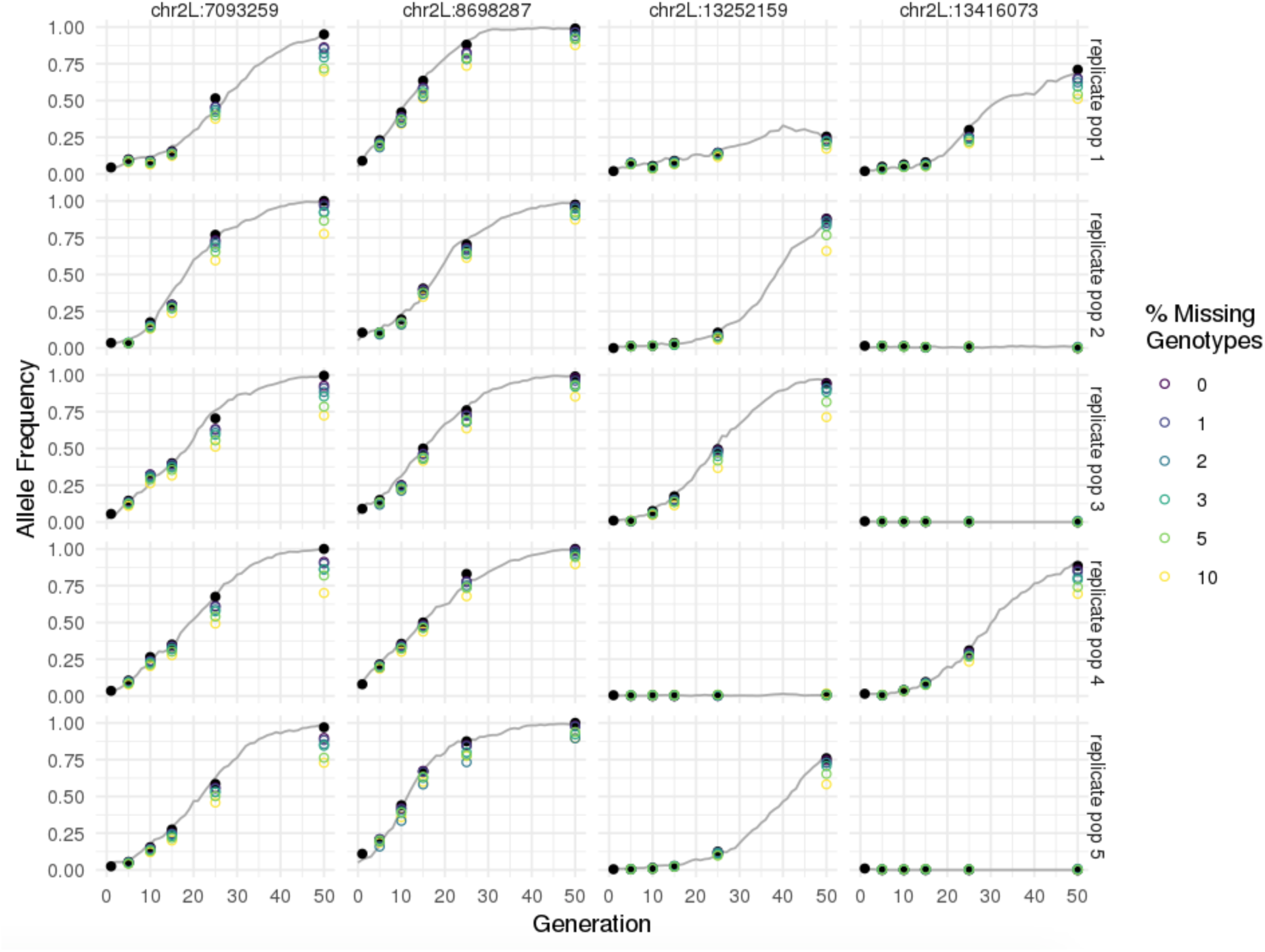
True population-wide allele frequencies (grey lines), true sampled chromosome allele frequencies (closed black circles) and HAFs (open circles) calculated at sites under selection (S=0.025) from samples simulated at 5x empirical coverage after 5,10,15,25, and 50 generations of recombination, using founder information with various fractions of missing of founder genotype calls (color).

**Supplemental Figure 7.**
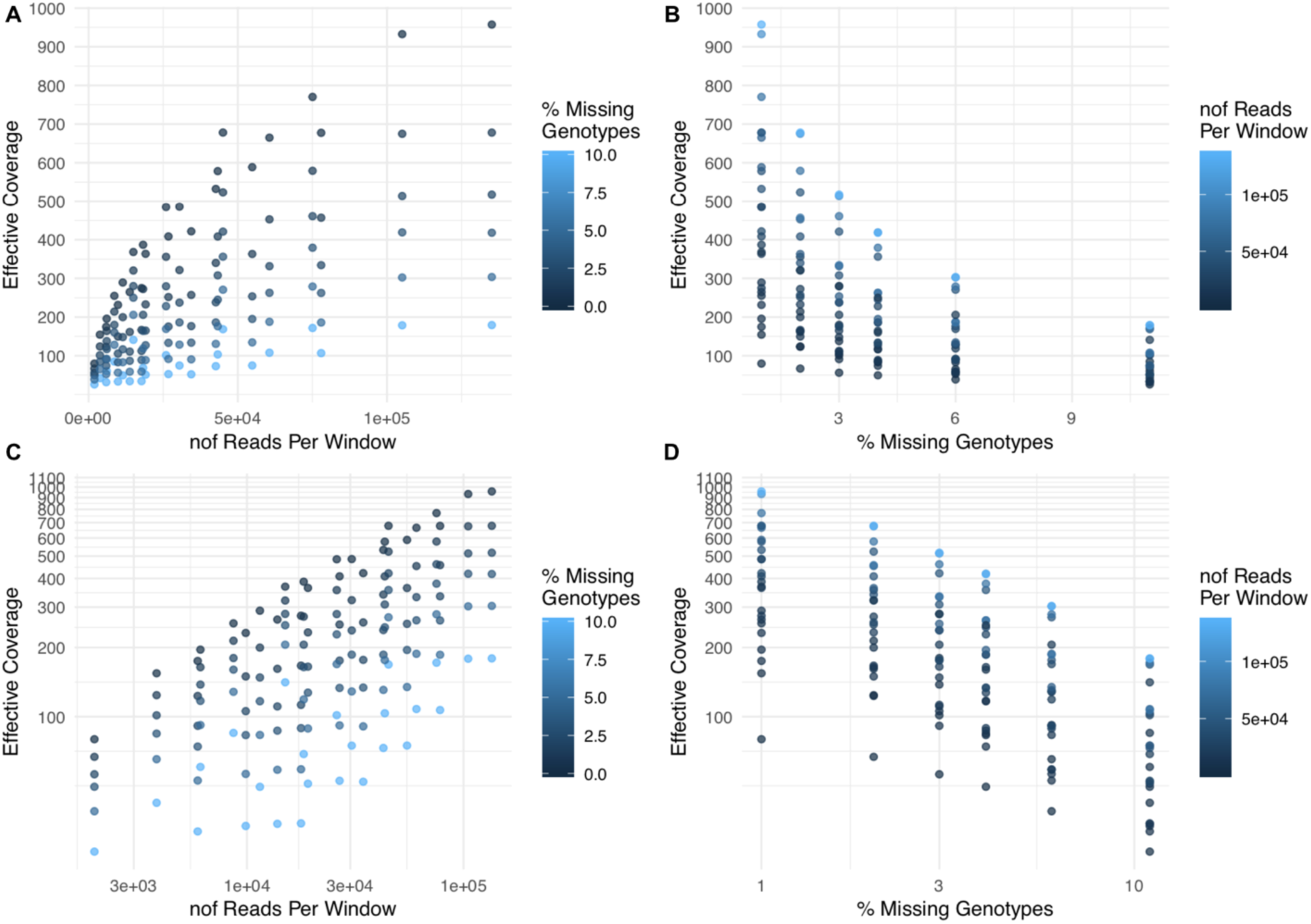
Relationship between effective coverage, number of reads per window, and percent of missing genotypes. he plots in the top row **(A-B)** indicate that the relationships are not linear. The plots in the bottom row **(C-D)** (where the x- and y-axes have been adjusted to log scale) suggest that the relationships are approximately log-linear.

